# A plastic vegetative growth threshold governs reproductive capacity in *Aspergillus nidulans*

**DOI:** 10.1101/050054

**Authors:** Luke M. Noble, Linda M. Holland, Alisha J. McLachlan, Alex Andrianopoulos

**Author notes:** Current address: Department of Biology, Center for Genomics and Systems Biology, New York University, NY 10012, USA. Current address: School of Biomolecular and Biomedical Science, Conway Institute, University College Dublin, Belfield, Dublin 4, Ireland (Linda M. Holland).

## Abstract

Threshold-limited ontogenic phases separating somatic growth from reproduction are a common feature of cellular life. Long recognized for flowering plants and animals, this life-history component may also be prevalent among multicellular fungi. We establish the environmental and genetic basis of developmental competence, the capacity to respond to induction of asexual development, in the model filamentous saprotroph *Aspergillus nidulans*. Density and pH are critical parameters for competence timing, and we identify five genes with heterochronic effects through genetic screens and candidate mutagenesis, including the conserved GTPase RasB and ambient pH sensor PalH. Inheritance of competence timing is quantitative, semi-dominant, transgressive, and extremely variable among progeny. Transcriptional profiling over competence acquisition demonstrates substantial activity in metabolic and signaling networks, highly concordant across species, and a wave of gene expression around centromeres indicative of chromatin remodeling. Competence, likely determined by species-specific endogenous hormones and metabolic capacity, governs much of biology associated with the mature fungal form – asexual and sexual reproduction, secondary metabolism, and, in some species, pathogenesis – and provides a new model for nutrient-limited life-history phases and their elaboration from unicellular origins.

## Introduction

Discrete states oriented toward growth or reproduction are fundamental to ontogenesis (Moran (1994), Istock and Istock (1967); Wilbur (1980)). In unicellular microbes they may be tightly integrated or even synonymous with the cell cycle (Johnston *et al.* (1977); Turner *et al.* (2012)), while in multicellular organisms intercellular signaling reflecting nutritional state is required to coordinate development. Plants pass through one or more vegetative phase changes before acquiring the capacity to flower (Poethig (2003); Poethig (2010)), many invertebrates attain the adult form through an intermediate juvenile stage (or stages) (Bishop *et al.* (2006); Truman and Riddiford (1999)), and mammals reach sexual maturity (Gluckman and Hanson (2006); Ahmed *et al.* (2009); Young (1976); Zacharias and Zacharias (1969)). All such examples involve interaction between the unfolding developmental program and nutrient accumulation, mediated by hormones.

Several genera of fungi, sister kingdom to animals within the Opistokhont supergroup, also require a minimum period of growth before acquisition of reproductive competence (Axelrod (1972); Noble and Andrianopoulos (2013)). While development of dormant, stress-resistant spores via asexual conidiation can be rapidly induced by exposure of a mature colony to an air interface and light, induction is without effect before a critical period of vegetative growth.

In contrast to plants and animals, competence in filamentous fungi is not associated with obvious morphological differentiation and, perhaps consequently, has received little attention since description in *Trichoderma* (Gressel *et al.* (1967)), *Penicillium* (Morton *et al.* (1958); Hadley and Harrold (1958)) and *Aspergillus* (Axelrod *et al.* (1973)) species (exceptions include Roncal and Ugalde (2003); Gravelat *et al.* (2008); Sheppard (2005); Ruger-Herreros *et al.* (2011); Adams *et al.* (1998); Nahlik *et al.* (2010)). Early work in *A. nidulans* showed competence acquisition is not simply a consequence of external nutrient limitation (Pastushok *et al.* (1976)), unlike analogous transitions in Dictyostelid amoebae and bacteria (Marin (1976); Bonner (2003)). Rapid, concerted changes in metabolic enzyme inducibility (Gealt and Axelrod (1974)), intracellular iron availability (Hall and Axelrod (1978)) and nutrient uptake rate were described, suggesting competence represents a transition between somewhat discrete organismal states (Kurtz and Champe (1979); Kurtz (1980)). More recently, competence for one species, *P. cyclopium*, was shown to be under control of a diterpenoid hormone (Roncal *et al.* (2002)) constitutively produced via the highly conserved mevalonate pathway, which produces diverse signaling compounds, such as steroids and insect juvenile hormones, through hydroxymethylglutaryl-CoA (Keller *et al.* (2005)).

While the genetic basis for synthesis and sensing of this cue remain unknown, regulation of growth and development by extracellular signals is known to be common in fungi, controlling spore germination inhibition, mating type recognition, negative autotropism, and metabolic influence on alternative reproductive strategies (Yu and Keller (2005); Vining (1990); Fox and Howlett (2008)). In the Aspergilli, much is also known of the molecular genetics of development, and of metabolism, hyphal extension, and mitosis during growth (Goldman and Osmani (2007)), but relatively little of organism level integration.

For *A. nidulans*, reproduction involves some combination of rapid vegetative growth, producing clonal nuclei distributed throughout the mycelium; asexual development, producing spores by the millions through iterated mitosis of stem-cell like phialides; parasexual development, resulting in chromosomal reassortment with rare recombination; and, given sufficient nutrients, sexual development within durable fruiting bodies (Figure 1A) (Pontecorvo (1952); Todd *et al.* (2007a); Champe *et al.* (1994); Etxebeste *et al.* (2010); Han (2009)). Differentiation is a response to conditions unfavorable to growth, such as the substrate-air interface. Asexual and sexual reproduction are reciprocally regulated by light during growth on solid substrate (Mooney and Yager (1990); Rodriguez-Romero *et al.* (2010)), but both are repressed indefinitely given sufficient nutrients in homogenous liquid culture. Upon establishment of a spatially patterned microenvironment during solid culture, developmental commitment is established in the colony interior through well-characterized regulatory networks. Upstream developmental activators (UDAs) – fungal specific transcription factors, conserved signal transduction components such as heterotrimeric G-proteins, and factors involved in production and perception of secreted signals – channel environmental cues into the late acting central developmental pathway (CDP), a positive feedback, morphogenetic network comprising transcription factors such as bristle (*brlA*) and abacus (*abaA*), which orchestrates conidiophore and asexual spore production (Clutterbuck (1969); Adams *et al.* (1988); Aguirre (1993); Sewall *et al.* (1990); Prade and Timberlake (1993); Andrianopoulos and Timberlake (1994)). Sexual development may follow, days later, and may be subject to a second competence threshold (Champe *et al.* (1981)).

**Figure 1.**
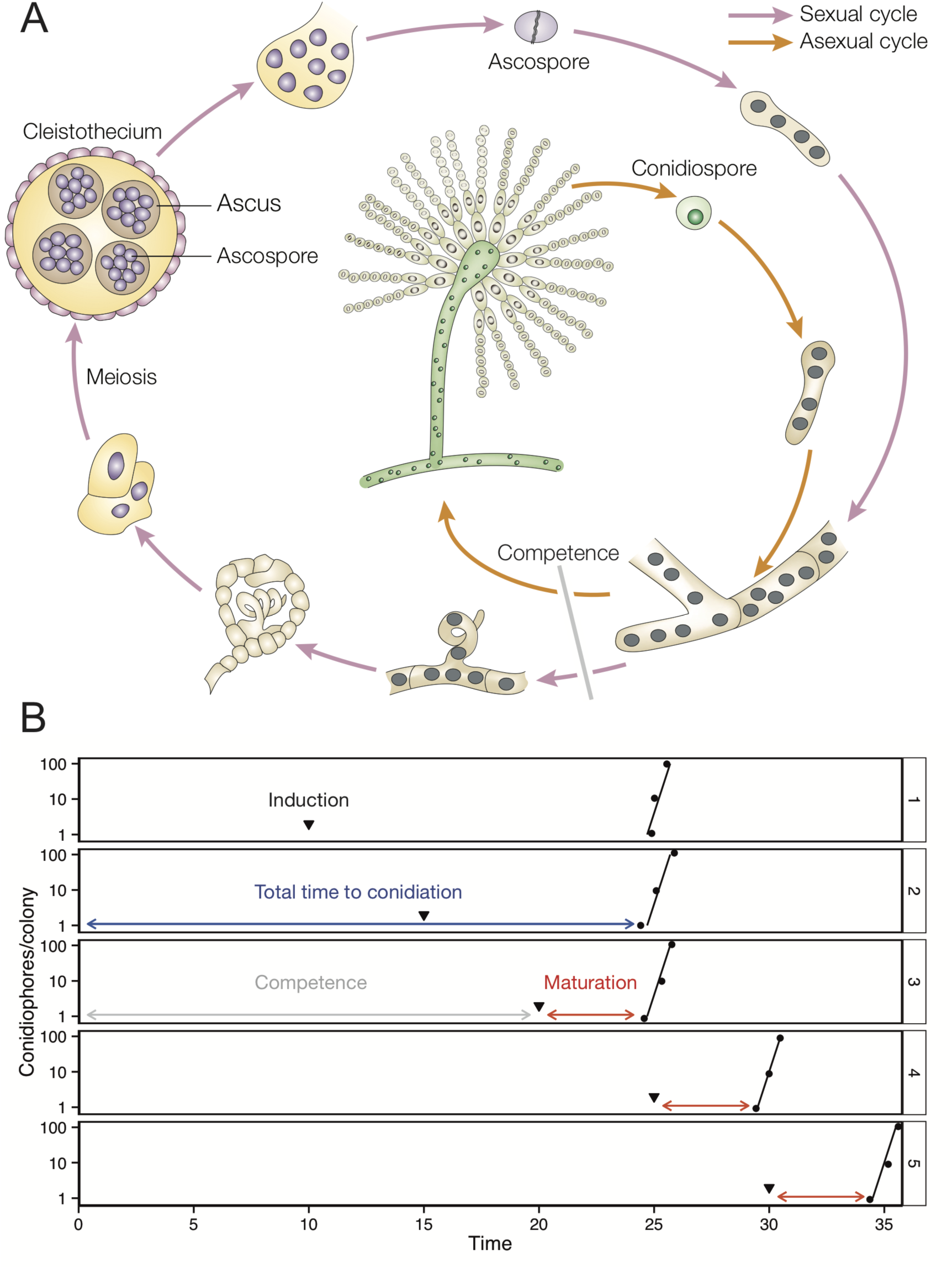
*A. nidulans* reproductive cycles and the logic of developmental competence. **A**. Three of the four reproductive cycles: asexual, sexual and hyphal growth, which may occur simultaneously in a single colony of sufficient age (adapted from Casselton and Zolan (2002)). In this work we focus on conidiation (asexual development), which is the first visible manifestation of developmental competence. Sexual development follows much later under conditions of sufficient nutrients and incubation in the dark. **B**. Defining competence in *A. nidulans*. Conidiophores per colony plotted by time since inoculation for five experiments varying the time of induction (black arrowheads) by exposure of single spore-derived colonies to air and light. Minimum time to conidiation (in blue) comprises the time required for competence acquisition (grey) and a constant maturation period for conidiophore production (red). Modified from Axelrod *et al.* (1973).

Early work on asexual developmental competence in *A. nidulans* yielded some genetic, as well as biochemical, insight. Precocious mutants were generated (Axelrod *et al.* (1973); Yager and Kurtz (1982); Butnick *et al.* (1984)), and cloning of temperature sensitive aconidial (*aco*) competence mutants identified subunits four and seven of the COP9 signalosome (CSN), with a third mapped to a small interval containing subunit five (Adams *et al.* (1992); Sutton (1999)). The CSN integrates nutrient sensing with the cell cycle in *S. cerevisiae* (Zemla *et al.* (2013)), and hormone sensing through regulation of proteasome activity in plants (Hind *et al.* (2011); Browse (2009)), but direct molecular connections to competence are lacking in *A. nidulans*. No genes underlying precocious competence have been identified and, more broadly, little is known of the role of environment, non-morphological aspects of differentiation such as polarization and transcriptional regulation, or the heritability of competence timing phenotypes.

## Results

### Environmental dependence of competence timing

We surveyed environmental and metabolic dependence of competence in a wild-type strain (WIM126) and two related, genetically undefined precocious strains. WIM27 (hereafter *prcA*) was isolated in a UV mutagenesis screen (Axelrod *et al.* (1973)) and WIM28 (hereafter *prcA; prcB*), with an enhanced phenotype, was generated by further mutagenesis of WIM27 (Kurtz and Champe (1979)).

Total time to conidiation consists of two components: competence, from breaking of spore dormancy to acquisition of reproductive capacity; and maturation, from induction of development to production of the conidiophore structure (Fig. 1A-B). Minimum total time to conidiation can be measured by growing single-spore derived colonies under inducing conditions (e.g. on the surface of an agar plate, under light). Maturation time can then be inferred by growing under non-inducing conditions (e.g. an agar plate overlaid with liquid medium) past this point, inducing development (e.g. by decanting the liquid overlay), and monitoring the time until conidiophore emergence.

Varying temperature, organism density, pH, primary metabolism and osmotic pressure, we found all but the last of these treatments altered competence timing significantly (Fig. 2A-D). We also found that competence and maturation often varied independently. As expected, competence was most rapid on primary carbon sources supporting rapid growth, such as glucose and fructose, and slowest on poorer, non-repressing sources such as glycerol and lactose (unpublished observations). On glucose medium, wild-type time to conidiation at 37° was 24.2 hours (± 0.07 hours standard error), including a maturation time of 4.1 ± 0.06 hours. Consistent with initial reports, *prcA* produced conidia around 2 hours earlier (22.6 ± 0.09), with *prcA; prcB* 1.5 hours earlier again (21.2 ± 0.05; Fig. 2A), and maturation timing was not significantly different to wild-type (*p* = 0.560 for *prcA*, *p* = 0.148 for *prcA; prcB*). The precocious phenotype was temperature dependent: the margin between wild-type and mutants, as a proportion of total time to competence, fell progressively with temperature such that the mutants were not significantly different to wild-type at 25°. For temperature, competence was most rapid within the optimal growth range (37-42°), as expected for metabolic dependence, while maturation was only slightly accelerated (ANOVA strain effect *p* < 2x10^−8^, temperature effect *p* < 10^−50^ for competence, 0.296 and 0.029 for maturation; Fig. 2A).

**Figure 2.**
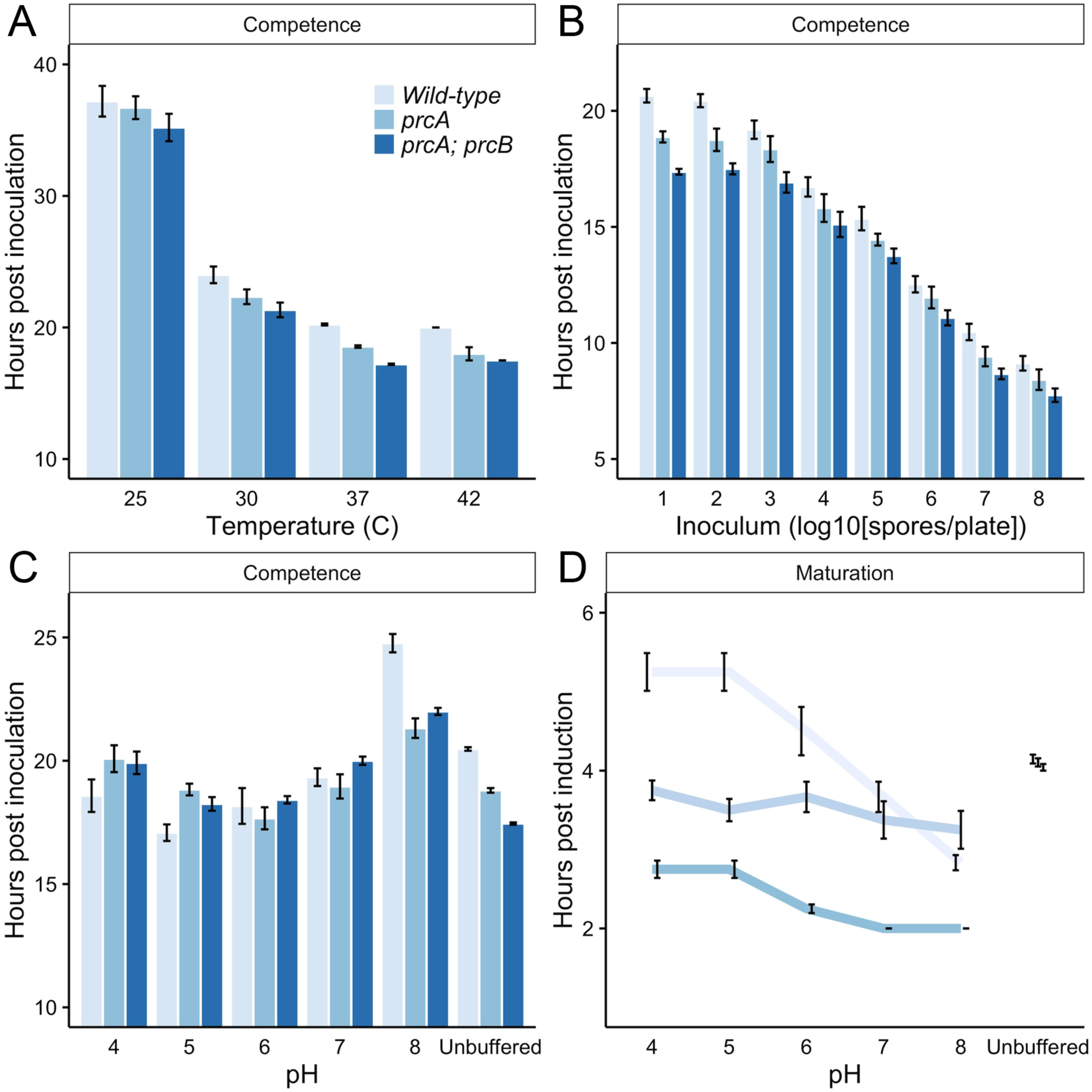
Environmental dependence of competence acquisition in wild-type and precocious mutants. **A**. Competence acquisition is temperature dependent, and the precocious phenotype is temperature sensitive. Mutants are significantly accelerated relative to wild-type (WIM27 *prcA p* = 2.25x10^−7^, WIM28 *prcA; prcB p* = 1.27x10^−13^ by linear model for 30-42°), but not at 25° (*prcA p* = 0.7, *prcA; prcB p* = 0.258). Maturation rate is not significantly different to that of wild-type: 4.1 ± 0.06 hours (*prcA p* = 0.927, *prcA; prcB p* = 0.121 by linear model). **B**. Competence acquisition is accelerated at densities above 10^3^ spores/plate (15.72 spores/cm^2^). *p* = 0.015 at 10^3^ spores/plate by linear model relative to 10 spores/plate. Maturation rate is not significantly different to wild-type (*prcA p* = 0.56, *prcA; prcB p* = 0.148 by linear model). **C, D**. Medium pH interacts with genotype and developmental stage to determine competence acquisition timing. Wild-type maturation rate is strongly pH dependent above pH 5. Maturation for the precocious mutants is reduced under all conditions, relative to unbuffered medium, and pH sensitivity is abolished (*prcA p* = 0.333) or dampened (*prcA; prcB p* = 0.0091). *prcA* and *prcA; prcB* show precocious competence for basic pH or unbuffered medium only, and *prcA; prcB* is significantly precocious relative to *prcA* only on unbuffered medium. Plotted data are mean ± standard error for at least three biological replicates per strain and condition.

Competence timing varied inversely with organism density for all strains, while maturation again remained relatively constant (strain effect *p* < 10^−13^, density effect *p* < 10^−70^ for competence, *prcA p* = 0.333 and *prcA; prcB p* = 0.096 for maturation; Fig. 2B). Effect size of the precocious phenotype was similar across the density range. Timing was identical for growth at an air interface on solid culture and for stationary liquid culture (unpublished observations), suggesting competence is not highly sensitive to colonial spatial organization or oxygen availability.

Buffering medium across a pH range significantly altered both competence and maturation timing in all strains, with effects dependent on both pH and genotype. Under acidic or neutral buffering (pH 7), the general effect on competence was to accelerate wild-type to similar, or even earlier, timing as the mutants (Fig. 2C). While wild-type at pH 6 was not significantly different to *prcA; prcB* on unbuffered medium, the mutant lines remained precocious under alkaline conditions, suggesting an acidity-mimicking alteration in ion homeostasis may underlie the precocious phenotype. Buffering at any pH removed entirely the phenotypic enhancement seen in *prcA; prcB* relative to *prcA*, however, further suggesting this strain carries a pH-sensitive mutation (or mutations), which can be phenocopied by external buffering. Variation in competence timing was driven largely by maturation, which varied near inversely across the pH range for wild-type (linear regression coefficient = −0.63, *p* = 0.0006), while the response was reduced for *prcA; prcB* (coefficient = −0.183, *p* = 0.0091) and not significant for *prcA* (coefficient = −0.11, *p* = 0.3333; Fig. 2D).

### Transcriptional profiling of competence acquisition

We profiled transcription during competence acquisition by microarray, first sampling wild-type mycelium hourly over a 5-hour spanning window (centered on competence at 14.5 hours for moderate density liquid culture) (Supplementary Fig. 1). At a false discovery rate of 5%, almost a quarter of the transcriptome was significantly differentially expressed at any single time-point. The global trajectory of gene expression over the window, relative to the earliest sample of 12 hours, was predominantly linear, with 1493 genes upregulated and 965 downregulated at 16 hours. This suggests competence is a transition between states, rather than a transient signalling event. The data were strongly correlated with published gene expression data from early vegetative growth of varying age in *A. nidulans* (Breakspear and Momany (2007)) and *A. fumigatus* (Sheppard (2005)), but not with germination in *A. nidulans* (Breakspear and Momany (2007)) or *A. niger* (Leeuwen *et al.* (2013)), showing these developmental landmarks are largely distinct (Fig. 3A). Overall concordance in expression direction between *A. nidulans* and *A. fumigatus* after competence acquisition was 74% (see Methods), representing almost one third of orthologous genes (Supplementary Fig. 2).

**Figure 3.**
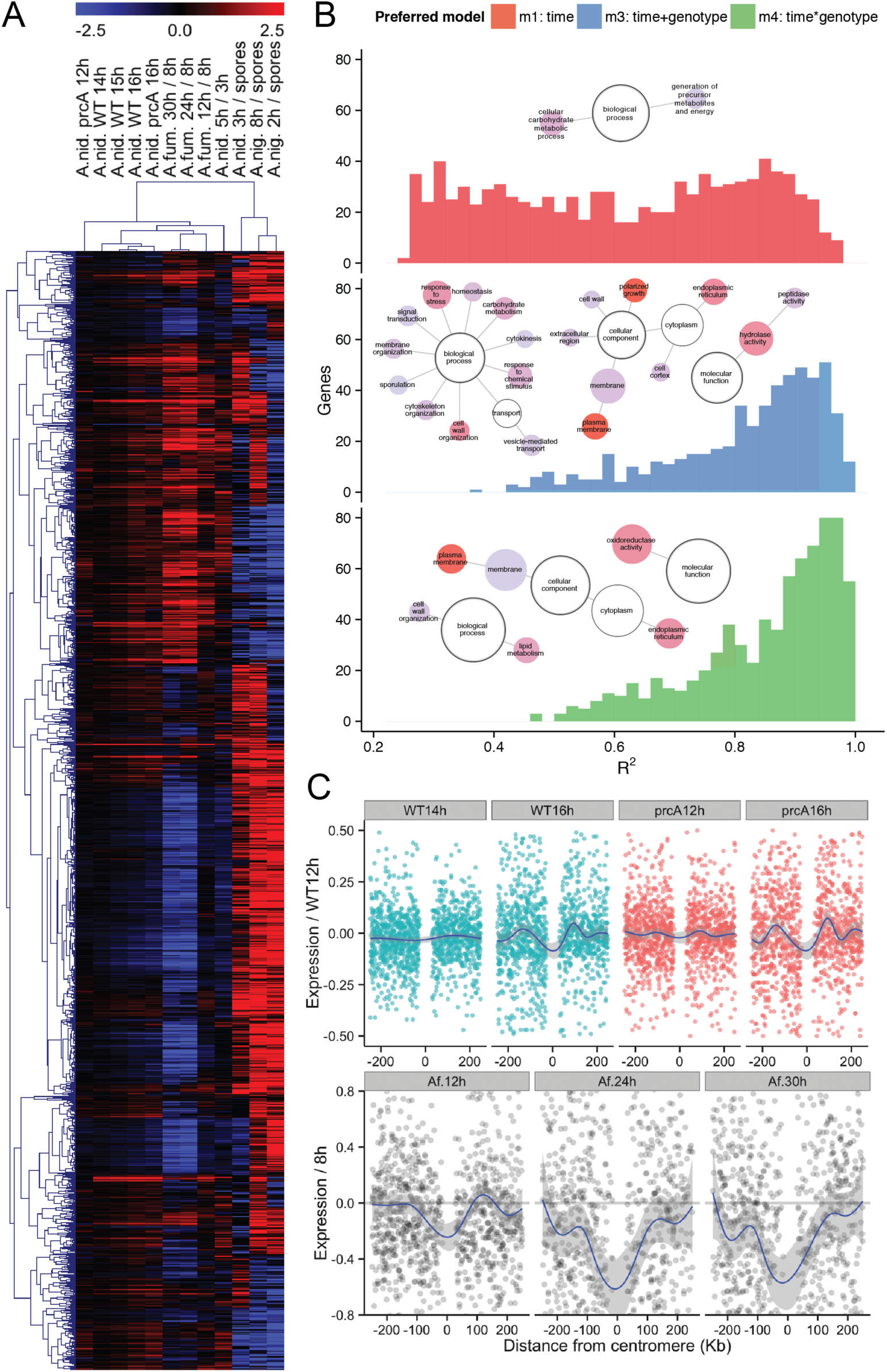
Transcriptional profiling of competence acquisition and comparative analysis. **A**. Hierarchical clustering of wild-type and *prcA* with *A. fumigatus* post-competence growth (12 vs. 8 h), *A. nidulans* during germination and early vegetative growth (3 h vs. spores, 5 h vs. 3 h), and *A. niger* during germination and early vegetative growth (2, 3 and 8 h vs. spores). Genes with missing data for more than two samples were excluded. Post-competence expression in wild-type *A. nidulans* is highly correlated with *prcA* (Spearmans *ρ* = 0.59 for wild-type 16 hours and *prcA* 16 hours), and with published data for *A. fumigatus* (*ρ* = 0.61 for wild-type 16 hours and *A. fumigatus* 12 hours) (Sheppard (2005)) and *A. nidulans* early vegetative growth (*ρ* = 0.59 for wild-type at 16 hours (here) and 5 hours from Breakspear and Momany (2007). **B**. Time and genotype-dependence of gene expression over competence acquisition. The distribution of correlation coefficients for each significantly differentially expressed gene is plotted for three of four linear models tested, where the model is preferred by Aikake Information Criterion (AIC; see Methods): time (model 1), genotype (m2; the smallest class, not shown), additive effects of time and genotype (m3) or additive and interaction effects of time and genotype (m4). Significantly enriched Gene Ontology (GO) terms among genes upregulated at competence are shown for each class, and for each significant GO hierarchy (central nodes in white). Node size represents the number of genes in each category, node color represents the significance of enrichment from white (no enrichment), through blue (*p* < 0.05 after multiple correction) and red (minimum of 1.672 x 10^−5^ for polarized growth under model 3) (Maere *et al.* (2005)). **C**. Epigenetic gene regulation in *A. nidulans* and *A. fumigatus* at competence acquisition. Gene expression relative to centromere position for wild-type and *prcA A. nidulans* (upper), and *A. fumigatus* (lower), with LOESS smoothing across all chromosomes. Competence acquisition is associated with establishment, or exaggeration, of a sinusoidal wave in expression symmetrical around the centromere, evident for *prcA* at 12 h (before competence at 12.5-13 h), and for wild-type *A. nidulans* after competence acquisition. Symmetry is reduced in *A. fumigatus* due to a more variable karyotype skewed toward submetacentric chromosomes.

To test for candidate gene expression differences that cause early competence acquisition, we next sampled the precocious mutant *prcA* over the window, testing for expression patterns best explained by (1) time, (2) genotype, (3) additive effects of both time and genotype, or (4) additive effects as well as interaction between these factors (Figure 3B). The mutant attains competence at 12.5-13 hours under the moderate density liquid culture conditions used. Functional analysis (Gene Ontology, Gene Set Enrichment Analysis, KEGG pathways) revealed highly significant activity: genes related to translation and ribosomal biogenesis (translation initiation and elongation, tRNA modification, RNA processing and transport) were progressively downregulated at and beyond competence acquisition, while transcripts annotated with signal transduction, polarized growth, endocytosis, plasma membrane modification, and vacuole and metal ion homeostasis functions increased in abundance (Fig. 3B, models 3 and 4). Dynamic regulation of metabolic genes was also apparent, spanning gluconeogenesis, ergosterol biosynthesis, the GABA shunt and central acetyl-CoA pathways, suggesting a shift from purely glycolytic primary metabolism, which may be maximal during early, pre-competence growth (Kurtz and Champe (1979); Kurtz (1980)). Accordingly, expression of the central energy homeostasis regulator pyruvate dehydrogenase kinase (PDK) increased steadily throughout the time course.

Gene expression during competence acquisition was also clearly altered at the level of chromosome structure or subnuclear localisation. First, competence acquisition was associated with establishment (or exaggeration) of a sinusoidal pattern in mean expression symmetrical around centromeres. This was accelerated in the precocious mutant, and marked in *A. fumigatus* (Fig. 3C). Second, expression from a number of verified and potential coregulated gene clusters was often altered in the precocious mutant, so that correlation with wild-type was often minimal at telomeres (Supplementary Fig. 3).

### Identification and genetic analysis of precocious mutants

Although a number of precocious competence mutants have been generated, underlying genes and the heritability of competence timing phenotypes are unknown. We undertook genetic and parasexual analysis of three undefined precocious mutant strains, two from the original UV mutagenesis screens (*prcA* and *prcA; prcB*) and a third (*prcC*) from a chemical mutagenesis screen (see Methods).

Mutants were first crossed to auxotrophic lines to introgress selectable markers, allowing diploid creation by complementation with a mapping strain marked on each of the eight chromosomes. Parasexual analysis, whereby haploids are regenerated with segregation of whole chromosomes, implicated loci of major effect on chromosome I for *prcB* and chromosome III for *prcA*, while results for *prcC* were ambiguous. Precocious competence timing in heterozygous diploids was, in all cases, dominant or semi-dominant to wild-type, however unambiguous designation was hampered by instability of the precocious phenotype through passaging and storage of spores. We note that instability of multiple precocious mutants was seen during Axelrod’s initial mutagenesis screens (Axelrod *et al.* (1973)), and we also observed this for two additional strains generated by repeating the screen.

Testing against a battery of environmental and chemical treatments revealed multiple phenotypes cosegregating with *prcB* in sexual crosses, including molybdate sensitivity and an inability to elicit contact inhibition from neighboring wild-type colonies (unpublished observations). Selection for restoration of molybdate tolerance by transformation with a genomic DNA library identified multiple unique inserts from a single locus on chromosome I that remediated both molybdate sensitivity and early competence acquisition, minimally spanning two genes (AN6885, AN6886). In parallel, we sequenced the genome of a *prcA; prcB* strain to 35X coverage (see Methods), detecting an early frameshift in AN6886, encoding the ambient pH sensor PalH. Molybdate sensitivity is a known phenotype of acidity-mimicking mutations, including palH loss of function mutations (Calcagno-Pizarelli *et al.* (2007); Negrete-Urtasun *et al.* (1999)). The frameshift cosegregated with molybdate sensitivity and the above phenotypes in eight F_1_ progeny by Sanger sequencing, and we conclude that *prcB* is a null allele of *palH*.

For *prcA* and *prcC*, pooled genomes for wild-type and precocious F_1_ backcross progeny were sequenced to a depth of approximately 160X each. For *prcC* we detected a frameshift in the *nsdD* gene, which encodes a GATA zinc finger transcription factor required for sexual development. A *nsdD*Δ deletion mutant showed precocious competence acquisition, consistent with a prior report of early conidiation for loss of function mutants (Han *et al.* (2001)).

From *prcA* and wild-type F_1_ pools, each comprising 12 haploid genomes, we detected 2153 single nucleotide variants (SNVs) and small insertion/deletions after filtering (see Methods). Most (1450) were fixed in both pools relative to the reference FGSCA4. Of 406 variants segregating and variable in frequency across pools (Supplementary Fig. 4), two SNVs were fixed in the precocious pools and at low frequency (1/12) in wild-type. Both were non-synonymous coding mutations with evidence for local linkage disequilibrium (Supplementary Fig. 5), located on chromosomes I and III. Single gene deletion mutants failed to alter developmental timing, however, and further parasexual analysis suggested the presence of chromosomal abnormalities (see Supplementary Data). Without resolution of the genetic basis of precocious competence in *prcA*, the possibility remained that phenotypic instability and quantitative inheritance seen in crosses of the progenitor (WIM27) and derived strains might be due to structural variation or multiple mutations.

### A mutational survey of candidate genes

We conducted candidate mutagenesis to study heritability for defined mutations in a standard genetic background, and broaden understanding of genetic influences on competence timing. First, we generated gain (G16V) and loss (T21N) of function point mutations altering activity of the *ras* family GTPase RasB, since activity of a related protein, RasA, has been shown to alter the timing of developmental transitions (although effects on competence were not shown; Som and Kolaparthi (1994)). In a *rasB*Δ deletion background, modulation of RasB activity also showed pleiotropic effects on growth and development. GTPase activity was anticorrelated with germination and conidiation (*p* < 10^−4^ and *p* < 10^−18^ by linear model, mutant genotypes coded as −1/1) while maturation rate was accelerated slightly by both alleles (ANOVA strain effect *p* < 10^−4^; Fig. 5A). The net effects on competence were strong in both directions: −1.63 ± 0.1 hours for RasB^T21N^ and 1.68 ± 0.18 hours for RasB^G16V^, Fig. 5B).

**Figure 4.**
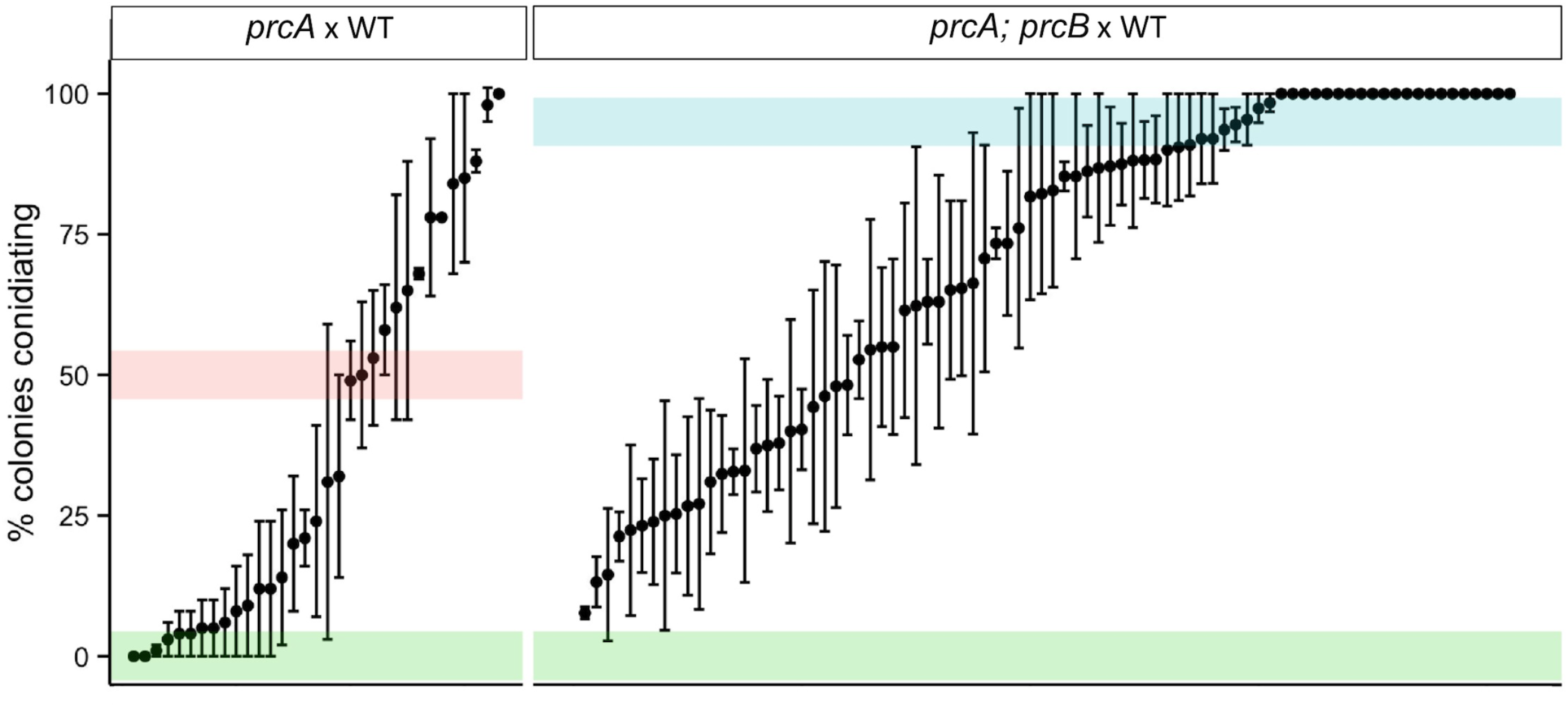
Competence timing is inherited as a continuous trait for precocious mutants. Conidiation timing (proportion of colonies conidiating after 22 hours growth) for a sample of F_1_ progeny from crosses between *prcA* and wild-type (WIM27 x WIM126, left) and *prcA; prcB* and wild-type (WIM28 x WIM126, right), mean ± standard error from 2-3 biological replicates. Shaded bars indicate parental timing (green for wild-type).

**Figure 5.**
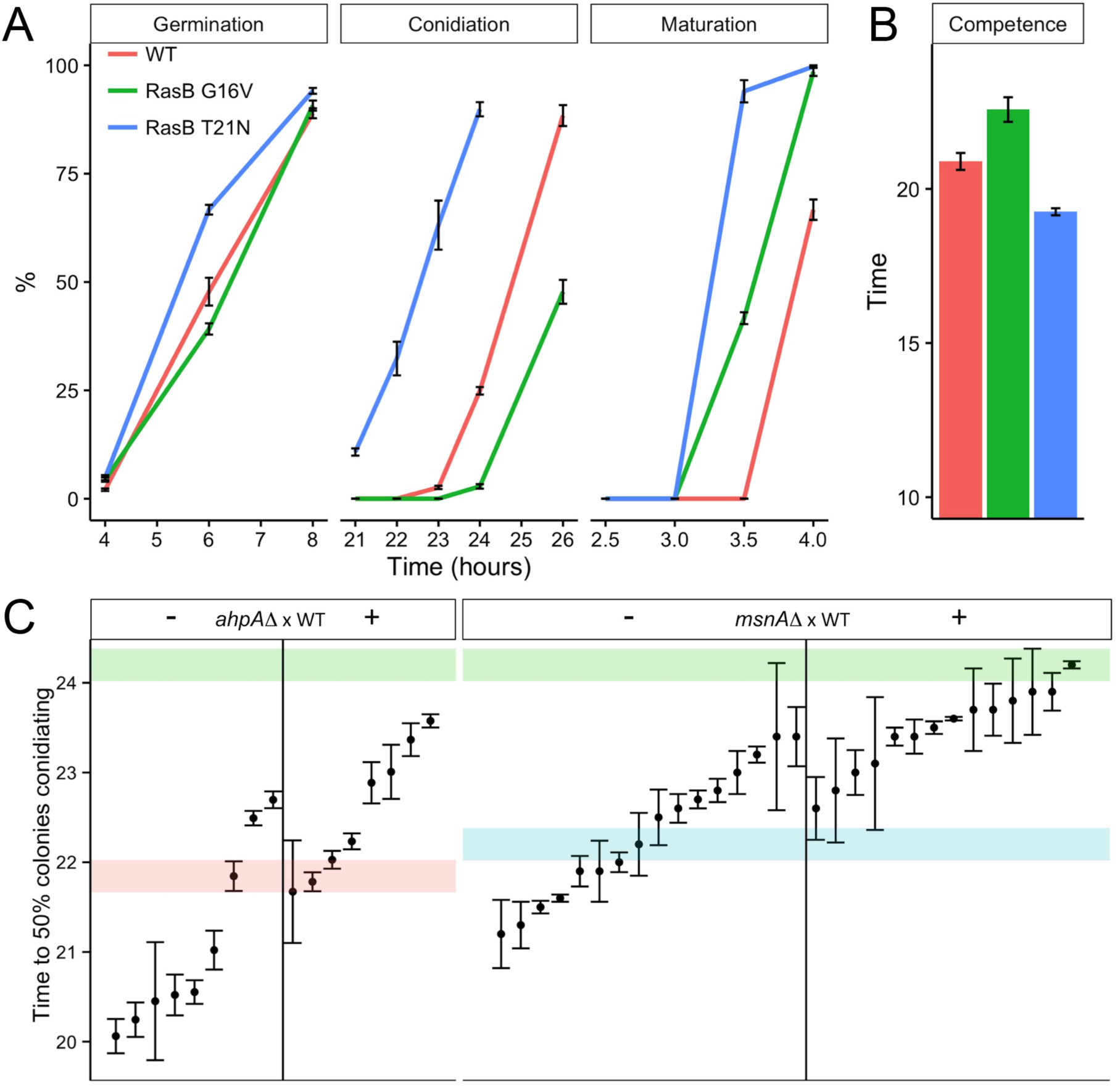
Genetic analysis of defined precocious mutants. **A**. RasB activity modulates the timing of multiple developmental landmarks. For gain (G16V) and loss (T21N) of function GTPase mutations, activity is negatively correlated with germination and conidiation timing, while both alleles slightly accelerate maturation timing. All strain contrasts are significant for germination at 6 h (*p* < 0.05 by two-sided Wilcoxon rank sum test), maturation at 3.5 h (*p* < 0.05), and for conidiation (*p* < 0.005 for a linear model including all time points, strain effect *p* < 10^−15^ by ANOVA). Data are mean percentages (± standard error) for at least three replicate experiments. **B**. Competence is significantly accelerated in RasB^T21N^ and delayed in RasB^G16V^ after accounting for differences in conidiation and maturation. Mean competence timing based on estimation of conidiation and maturation timing (50% colonies conidiating) by linear regression (± additive standard errors). **C**. Competence timing for the F_1_ of backcrosses of precocious deletion mutants *ahpA*Δ and *msnA*Δ to their wild-type transformation strains. In both cases, F_1_ timing is shifted toward that of the precocious parent, mean timing by genotype is significantly different (*p* = 5.2x10^−6^ and 1.3x10^−4^, by Wilcoxon rank sum test), and progeny transgress the precocious parent in range. Data are mean ± standard error of 3 biological replicates.

We additionally generated and analyzed 70 targeted deletion or disruption mutants (see Methods), selected on criteria of differential expression over competence acquisition, predicted gene function (predominantly transcription factors) and evolutionary conservation. Null mutants of two genes showed significantly earlier competence acquisition: homeodomain transcription factor *ahpA* (21.84 ± 0.165 hours), and C_2_H_2_ transcription factor *msnA* (22.2 ± 0.35 hours; see Supplementary Data). Backcrossing of both mutant strains showed transgressive, continuous inheritance of competence timing, as seen for *prcA* strains (Fig. 6B). In both cases, precocious competence was partially dominant in the F_1_ – for *ahpA*Δ, no progeny of wild-type timing were seen – and additionally dependent on F_1_ genotype (*p* = 5.2x10^−6^ for *ahpA*Δ, 1.3x10^−4^ for *msnA*Δ, two-sided Wilcoxon rank sum test). Quantitative and highly variable heritability of competence timing is therefore inherent to the precocious phenotype, not an artifact of background variation segregating in *prcA* strains.

## Discussion

We have outlined the environmental, genetic, and transcriptional basis of a reproductive maturity life-history threshold in *A. nidulans*, conserved throughout filamentous Ascomycetes at least. Competence varies with temperature, pH and quality of carbon source, consistent with an interaction between primary metabolism and a density-dependent signal. We have focused on asexual development, as the earliest manifestation of competence and by far the better studied reproductive mode. An additional, much later threshold for competence to respond to sexual development has also been demonstrated using a temperature sensitive allele of the pivotal regulator *velvetA* (Champe *et al.* (1981)).

### Transcriptional and colonial differentiation during development

Molecular genetic studies have shown the expression of many genes changes, in quantity or localization, during early vegetative growth (Garzia *et al.* (2009); Etxebeste *et al.* (2009); Etxebeste *et al.* (2008); Miller *et al.* (1991); Kwon *et al.* (2010); Soid-Raggi *et al.* (2006); Harispe *et al.* (2008); Kawasaki and Aguirre (2001); Tsitsigiannis (2004)), and biochemical and cell biological studies have demonstrated some broad effects of age on cellular state (Gealt and Axelrod (1974); Hall and Axelrod (1977)). Competence timing has rarely been determined in these studies, however, and so the relationship between developmental and chronological age has been confounded. We find that while differentiation during competence acquisition is not apparent at the level of morphology or growth kinetics, transcriptional differentiation is extensive and highly concordant across the two Aspergilli examined.

While multiple physiological correlates of competence acquisition show non-linear, switch-like dynamics, we observed a predominantly monotonic transition in gene expression over the 4-hour window suggesting a reduction in biosynthetic activity and an increase in signaling, development and secondary metabolic gene networks. Altered asymmetry at the level of the colony might be a trivial explanation of some of the broad patterns of gene expression observed. Kinetics and morphology of early filamentous growth – germination, nuclear division, septation and branching – have been studied extensively (Fiddy *et al.* (1976); Dynesen and Nielsen (2003); Roper *et al.* (2011); Rittenour *et al.* (2009); Trinci (1974)), but over a wide range of conditions and rarely with regard to competence state, complicating inference. Dynesen and Nielsen monitored the hyphal growth unit (the ratio between total hyphal length and the number of hyphal tips) in low-density stationary cultures at 30° and found it approaches a constant value soon after hyphae possess 8-10 tips (around 14 hours in rich medium) (Dynesen and Nielsen (2003)). Based on the range of growth conditions examined here, this would be well before competence acquisition, ruling out a change in apical to subapical compartment ratio. Asymmetry in metabolic and transcriptional activity within the colony, for which there is abundant evidence at diverse stages of vegetative growth, may contribute to, or even drive, the observed patterns, however.

Documented biochemical changes in hyphal differentiation from unpatterned to polar growth include establishment of apical-high gradients of reactive oxygen species, Ca^2+^ and protons (Robson *et al.* (1996); Rodriguez-Urra *et al.* (2012); Roncal *et al.* (1993); Semighini and Harris (2008)). It seems likely, upon case-by-case examination of experimental conditions that these aspects of hyphal differentiation precede competence acquisition. Importantly, however, the FlbB/FlbE regulatory complex appears to assemble apically at or shortly after competence acquisition, linking developmental induction capacity with the morphogenetic, positive feedback, central development pathway (Clutterbuck (1969); Adams *et al.* (1988); Aguirre (1993); Sewall *et al.* (1990)). It remains to establish the precise timing of these and other hyphal differentiation events with respect to density and competence, and the potential for spatial regulation of transcription.

Functional analysis of gene expression over the competence acquisition window supports early biochemical studies showing a progressive decrease in rate of transport of glucose and other metabolites (Kurtz and Champe (1979); Kurtz (1980)), and a sharp change in iron availability or oxidative state (Hall and Axelrod (1978)). Strong enrichments were seen among upregulated genes for primary metabolism and other iron intensive pathways such as ergosterol biosynthesis, associated with an oxidative stress response. Downregulated activity was mainly related to ribosome biogenesis and translation, and appeared to follow the stress response. Correlation between mRNA and protein abundance is generally good, where it has been tested, in fungi (*r* 0.4-0.7) (Lee *et al.* (2011); Vogel and Marcotte (2012); Vogel *et al.* (2011)), but may be weaker for downregulated genes and for specific classes such as ribosomal biogenesis, since these proteins may be relatively stable. Such systemic changes are apparently without consequence for growth rate, which is exponential for several hours before and after competence acquisition (Hall and Axelrod (1978)). In all, the data support the view that the precompetent growth phase is dedicated principally to rapid biomass accumulation, and that competence acquisition elicits a significant stress response, with redistribution of reducing power to secondary metabolic pathways and alterations to intracellular, and perhaps extracellular, transport.

### Genetic and epigenetic basis of competence timing

Instability of multiple precocious mutants was seen during Axelrod’s initial mutagenesis screens (Axelrod *et al.* (1973)), and we again observed this for two additional strains generated by repeating the screen. This suggested the possibility some precocious strains might not be true genetic mutants, with variability in competence timing allowing selection of transiently heritable epigenetic mutants. Alternatively, if competence is genetically robust, the effect of mutational perturbation might be compensated over time.

Genetic analysis of *prcA* and *prcA; prcB* – where precocious competence has been stably maintained almost 40 years since isolation – showed competence timing was inherited as a continuous quantitative trait with variable stability among progeny. F_1_ progeny scored as precocious immediately after isolation often reverted to wild-type timing after storage (wild-type spores are stable for months in water at 4°), while others remained stably precocious under identical conditions. This suggested only a subset of initially precocious progeny might carry underlying genetic variation, and that biochemical activity in resting spores might reset competence timing. Less ambiguously, progeny of defined precocious mutants *ahpA*Δ and *msnA*Δ also showed continuous, transgressive variation in competence timing, with the F_1_ genotypic classes overlapping in phenotypic values. This demonstrates that timing is a considerably plastic trait, at least as assayed here over a single generation. Monitoring over additional generations is required to additionally address the possibility of the robustness of wild-type timing to genetic perturbation.

Partial dominance in the F_1_ suggests shared cellular environment during meiosis exerts an effect on ascospore composition, which is maintained through vegetative growth and transmitted to the asexually derived conidia used for assay. The long-term stability of this transgenerational inheritance is unknown, and many other unanswered questions remain, not least the basis of variable penetrance. Small RNAs have been implicated in phenotypic plasticity in other species (e.g. Calo *et al.* (2014)) and, as in analogous life-history transitions in other systems, parental nutrient deposition (e.g. linolenic fatty acids in asexual spores (Vanetten and Gottlieb (1965); Evans and Gealt (1985))) may influence competence timing and the observed mode of inheritance.

### Evolution of reproductive competence and complex life cycles

Differentiation is typically considered as the generation of discrete cell types, and life-history stages have historically been delineated as simple (direct) or complex (indirect) (Moran (1994); Wiegmann *et al.* (2009)). One definition of this complexity is an abrupt ontogenetic change in an individual’s morphology, physiology, and behavior, usually associated with a change in habitat (Wilbur (1980)), while others have elevated ecological or morphological criteria (Istock and Istock (1967)). Clearly, a narrow definition is more broadly defendable, as physiological differentiation is likely to be pervasive, particularly in organisms of limited morphological diversity (Beadle *et al.* (1938); Stoodley *et al.* (2002); Agathocleous and Harris (2013)). Still, competence in filamentous fungi appears to be broadly analogous to nutrient-limited, hormone-mediated early life-history stages in animals and plants. Minimum time to competence occurs within temperature and pH ranges optimal for growth, on preferred carbon and nitrogen sources (according to hierarchical catabolite repression systems (Hynes and Kelly (1977); Wong *et al.* (2008))). Competence timing is density dependent, presumably as a response to resource colonization (the response to other species, or potentially even heterokaryon incompatible strains of the same species, would be expected to differ), and a diterpenoid hormone sufficient for both competence acquisition and induction of asexual development has been isolated from *P. cyclopium* (Roncal *et al.* (2002); Roncal and Ugalde (2003)).

Ultimately, development and primary metabolism must be temporally or spatially segregated activities (Adams *et al.* (1988); Adams and Timberlake (1990)). For filamentous fungi, this may be ensured by delayed reproductive capacity and the mycelial growth habit, with exploratory nutrient acquisition at the colony margins and a depleted interior committed to secondary metabolism and reproduction. Beyond this basic necessity, the precise timing of competence acquisition presumably reflects life-history optimization for colonization of ephemeral, patchy environments (Andrews (1992); Heaton *et al.* (2016)). A bang-bang strategy – complete allocation toward vegetative growth, followed by a resource-dependent switch to complete allocation toward spore production – has been predicted to be optimal for an asexual saprobe (Gilchrist *et al.* (2006)). Under this patch array model, fitness is maximized by efficient colonization of newly arising patches of fixed resource levels, which involves trade-offs between spore production schedule and quantity, as well as propagule quantity and quality (motility, durability). A genetically assimilated competence schedule should then reflect the species’ present and historical ecology, and it’s level of predictability, particularly for critical factors such as patch size, quality and density. Variance in patch size and distribution (Gilchrist *et al.* (2006)), as well as advantages accruing from later, more costly sexual development, may explain the apparent superiority of a mixed strategy of continued vegetative growth, along with environmentally induced asexual development, over the finality of a post-competence reproductive bang. Diurnal cycles of light, temperature and air movement, of obvious relevance to patch availability and spore dispersal, may also contribute to the observed set point of competence + maturation at 24 hours, although it remains to see if this holds outside laboratory conditions, and for species other than *A. nidulans*.

The existence of a threshold-limited life-history trait in diverse fungi of relatively simple morphological complexity, coupled with increasingly dense phylogenetic sampling and genomic resources in a number of clades, offers a number of interesting opportunities. For one, our understanding of the underlying metabolic basis of more familiar adult fungal activities would gain from studying how regulatory controls are imposed at the colony level on a precompetent state committed solely to rapid cell division and hyphal growth. More broadly, further study of competence timing, and its variation within and across species, would allow testable inferences of fungal ecology and life-history theory. Lastly, although nutrient limited thresholds are ubiquitous in complex life cycles, their bioenergetic basis is in most cases poorly understood. Comparative study of the genetic basis of competence across fungal phylogeny would yield insight into the integration of conserved growth and divergent developmental networks, and of the evolution of complex life cycles from simple beginnings.

## Materials and Methods

### Strains and culture conditions

Strains used during this study are shown in Supplementary Table 1. All are derived from the reference Glasgow isolate FGSCA4 (Arnaud *et al.* (2009)). Minimal medium (ANM + 10 mM ((NH_4_)_2_T), carbon free (CF), complete medium (CM) and basic growth conditions, handling, genetic and parasexual analysis were as described (Todd *et al.* (2007b); Todd *et al.* (2007a)). Yeast extract glucose (YEG) medium contained 2% glucose and 0.5% yeast extract. Alternative carbon sources were added to CF medium to 1% w/v. Assays to detect molybdate sensitivity were conducted as described (Arst *et al.* (1970)). Buffering of 2% high-grade agar minimal medium used citrate/sodium citrate (pH 4-6) (Gomori (1946)) and phosphate/citrate (pH 6-8) (Pearse (1991)) buffers at a final concentration of 10 mM.

### Competence timing assays

All assays were performed at low density on solid culture (<=100 colonies per 90 mm diameter plate), using spore dilutions quantified by hemocytometer and stored at 4° in H_2_0 for up to one month. Testing of specific pH, metabolic substrates or chemical agents was on minimal medium, all other assays used unbuffered complete medium.

Competence acquisition is the total time to conidiation, less maturation (time from induction to visible production of conidiophore vesicles). Conidiation was defined as the mean time, from inoculation, at which 50% of at least 30 isolated colonies had produced a conidiophore vesicle, for at least three replicate plates. Maturation time was defined identically except time zero was induction of competent colonies, achieved by decanting a 15 ml liquid medium overlay. Plates were incubated at 37° under constant fluorescent white light (1 m from polycarbonate shielded F36W/840 triphosphor fluorescent white lighting, Sylvania Lamps).

### Generation of gene deletion, disruption and transgenic strains

Transformation strains carried an *nkuA* deletion to maximize the frequency of homologous integration (Nayak (2005)). For gene disruption, PCR primers targeted to the 5’ coding region of genes were designed with an amplicon size of 400-600 bp. Products were cloned (plasmid Af pyro Pst H3 EI SK+ (*Ssp*I)) for transformation of strain TN02A25, selecting for pyridoxine prototrophy. Putative transformants were screened for the presence of the transforming plasmid by PCR against *A. fumigatus pyroA*. For genetic analysis of strains with a competence phenotype, the parental transformation strain was restored to riboflavin prototrophy to allow efficient backcrossing. Segregating auxotrophic markers, which do not alter competence timing on supplemented medium, were scored in crosses to confirm Mendelian segregation.

Deletion of *rasB* was by gene replacement in MUS51BARB12 with the *Neurospora crassa pyr-4* marker, site-directed mutagenesis was by inverse PCR and confirmed by Sanger sequencing. GTPase mutants were targeted to the *yA* locus in MUS51BARB12 and assayed in a *rasB*Δ background. For all other targeted gene deletions, cassettes based on the *A. fumigatus pyrG* selectable marker were obtained from the FGSC and amplified by PCR for transformation of strains PPU+ and MUS51BARB12. Gene replacement is highly efficient in *A. nidulans nkuA* strains (Nayak (2005)), and 5-100 transformants were typically obtained for each transformation. The AMA1 genomic library used for suppression of *prcB* molybdate sensivity has been described previously (Aleksenko and Clutterbuck (1997)).

### Mutagenesis screens

A screen for precocious competence acquisition followed Axelrod (1973), with mutagenesis by treatment with 4-nitroquinoline oxide to 70% lethality. Generation of the precocious strain PCV9 (*prcC*) used a filtration enrichment screen based on an unpublished observation of interaction between competence state and *velvetA* (*veA*) genotype (Kennedy (1999)). As reported for a *veA*_*1*_ mutant, ectopic expression of *brlA* drives conidiation in liquid culture (Adams *et al.* (1988)). This holds for *veA*_*1*_ and *veA^+^* in the incompetent state, but not for competent *veA^+^*. The ability to recover conidia from liquid cultured mycelium produced by *brlA* overexpression in a wild-type background therefore reflects competence state. A *veA^+^* strain carrying *brlA* under control of the inducible *alcA* promoter (pTA29, Adams *et al.* (1988)) was mutagenized as above, grown to competence in glucose minimal medium, then switched to 20 mM threonine to induce *brlA* expression for 12 hours. Production of conidia is expected from (1) mutants that have not yet acquired competence or are incapable of doing so, and (2) mutants that mimic *veA*_*1*_ or otherwise suppress *veA^+^*. PCV9 fell into the latter class, and was subsequently found to be precocious.

### Genome sequencing

Genome sequencing for a single *prcA; prcB* strain (PABH13) was performed at the Allan Wilson Genome Centre (75 bp single end reads, Illumina Genome Analyser II). Sequencing of mutant and wild-type bulked segregant pools for *prcA* (12 F_1_ progeny), *prcA; prcB* (12 F_1_ progeny) and *prcC* (4 F_1_ progeny) was by Macrogen (100 bp paired-end reads, Illumina HiSeq 2000). Sequence data are available from NCBI SRA (BioProject PRJNA319684).

For reference-based variant calling, reads were mapped by BWA mem (0.7.3) (Li and Durbin (2010)) with default settings to *A. nidulans* FGSC4 reference genome version s04-m05-r02 (AspGD) and processed with samtools (0.1.16) (Li *et al.* (2009)) and Picard (http://picard.sourceforge.net) utilities. Variants were called using the Genome Analysis Toolkit (GATK) HaplotypeCaller (2.6-4) for haploid samples or UnifiedGenotyper for pooled samples, and filtered by mapping quality (MQ < 40), depth (10 < DP < 2X mean coverage), strand bias (FS > 60), variant position relative to reads (ReadPosRankSum < −8), and variant quality/depth (QD < 2) (McKenna *et al.* (2010); Li (2011)). Remaining variants within regions masked by RepeatMasker (4.03) or clustered around centromeric gaps in the reference sequence were also excluded. For bulked sequencing, singleton calls across combined pools were removed. Functional predictions were generated by snpEff (3.0) (Cingolani *et al.* (2014)). Structural variation was analyzed with BreakDancer (1.3.6) (Chen *et al.* (2009)) and pindel (0.2.4) (Ye *et al.* (2009)), and by genome alignment after de novo assembly. Assembly used ABySS (1.3.4) (Simpson *et al.* (2009)) after Quake (0.3.2) read correction (Kelley *et al.* (2010)). Contigs were aligned to the reference with MUMmer (3.22) (Kurtz *et al.* (2004)) or by the UCSC chain/net pipeline (Kent *et al.* (2011)).

### Microarrays, data processing and analysis

Mycelium for wild-type and *prcA* microarrays spanning competence acquisition were from 145 mm diameter petri dishes containing 50 ml YEG inoculated with 10^5^ conidia ml^−1^, incubated at 37°. Between 10 and 50 plates were pooled per time point, harvested by vacuum filtration and immediately frozen in liquid nitrogen. Total RNA was prepared in TRIzol (Invitrogen) after disruption by 1 mm zirconia beads (Daintree Scientific) in a FastPrep FP120 (MPBio). Poly-A RNA was isolated on oligo-(dT) cellulose (Sigma), reverse transcribed using the Invitrogen Superscript Indirect Labeling Kit with random primers, and cDNA purified using Zymo DNA Clean & Concentrator columns according to Pathogen Functional Genomics Resource Center (PFGRC) protocol M007 (http://pfgrc.jcvi.org/index.php/microarray/). Samples were quantified by NanoDrop and competitively hybridized to version 2 whole genome glass slide microarrays provided by the PFGRC, according to protocol M008.

For wild-type a closed-loop circuit with a minimum of three dye-swapped competitive hybridizations for each time point was used, for a total of 16 hybridizations (see Supplementary Fig. 1). For *prcA*, two biological replicates as dye-swaps were hybridized per time point against wild-type, for a total of six hybridizations. Microarrays were scanned on an Axon GenePix 4100 or 4000B, ensuring channel ratios were in the range 0.95-1.05. Raw data were processed using the R package (http://www.r-project.org/) limma, with background correction (normexp offset = 50) and duplicateCorrelation for replicate probes, normalized to wild-type at 12 hours (Ritchie *et al.* (2007); Oshlack *et al.* (2007); Smyth (2004); Smyth and Speed (2003); Smyth *et al.* (2005)). Raw and processed data are available from NCBI GEO (GSE68352).

## Acknowledgements

We thank Mary Teo, Katrin Hubner and Sophie Davidson for experimental contributions, Len Kelly, Phil Batterham and members of the Hynes/Davis/Andrianopoulos laboratory for constructive discussions, Alicia Oshlack for statistical advice, and Tom Harrop, Annalise Paaby, Christina Zakas, Joe Heitman and, especially, David Axelrod for improving the manuscript. This work was supported by a grant from the Australian Research Council to AA and an Australian Postgraduate Award to LMN. LMH is supported by the Irish Research Council ELEVATE International Career Development fellowship-cofunded by Marie Curie Actions.

## References

Adams T. H., Boylan M. T., Timberlake W. E., 1988 brlA is necessary and sufficient to direct conidiophore development in Aspergillus nidulans. Cell 54: 353–362.

Adams T. H., Timberlake W. E., 1990 Developmental repression of growth and gene expression in Aspergillus. Proceedings of the National Academy of Sciences of the United States of America 87: 5405–5409.

Adams T. H., Hide W. A., Hide W. A., Yager L. N., Lee B. N., 1992 Isolation of a gene required for programmed initiation of development by Aspergillus nidulans. Molecular and Cellular Biology 12: 3827–3833.

Adams T. H., Wieser J. K., Wieser J. K., Yu J. H., 1998 Asexual sporulation in Aspergillus nidulans. Microbiology and Molecular Biology Reviews 62: 35–54.

Agathocleous M., Harris W. A., 2013 Metabolism in physiological cell proliferation and differentiation. Trends in Cell Biology 23: 484–492.

Aguirre J., 1993 Spatial and temporal controls of the Aspergillus brlA developmental regulatory gene. Molecular Microbiology 8: 211–218.

Ahmed M. L., Ong K. K., Dunger D. B., 2009 Childhood obesity and the timing of puberty. Trends in Endocrinology & Metabolism 20: 237–242.

Aleksenko A., Clutterbuck A. J., 1997 Autonomous Plasmid Replication inAspergillus nidulans:AMA1 and MATE Elements. Fungal Genetics and Biology 21: 373–387.

Andrews J. H., 1992 Fungal life-history strategies. In: Carroll GC, Wicklow DT (Eds.), The fungal community: Its organization and role in the ecosystem, second edition, CRC Press, p. 952.

Andrianopoulos A., Timberlake W. E., 1994 The Aspergillus nidulans abaA gene encodes a transcriptional activator that acts as a genetic switch to control development. Molecular and Cellular Biology 14: 2503–2515.

Arnaud M. B., Chibucos M. C., Costanzo M. C., Crabtree J., Inglis D. O., Lotia A., Orvis J., Shah P., Skrzypek M. S., Binkley G., Miyasato S. R., Wortman J. R., Sherlock G., 2009 The Aspergillus Genome Database, a curated comparative genomics resource for gene, protein and sequence information for the Aspergillus research community. Nucleic Acids Research 38: D420–D427.

Arst H. N., MacDonald D. W., MacDonald D. W., Cove D. J., 1970 Molybdate metabolism in Aspergillus nidulans. I. Mutations affecting nitrate reductase and-or xanthine dehydrogenase. Molecular & general genetics: MGG 108: 129–145.

Axelrod D. E., 1972 Kinetics of differentiation of conidiophores and conidia by colonies of Aspergillus nidulans. Journal of general microbiology 73: 181–184.

Axelrod D. E., Gealt M., Gealt M., Pastushok M., 1973 Gene control of developmental competence in Aspergillus nidulans. Developmental Biology 34: 9–15.

Beadle G. W., Tatum E. L., Clancy C. W., 1938 Food level in relation to rate of development and eye pigmentation in Drosophila melanogaster. The Biological bulletin 75: 447–462.

Bishop C. D., Huggett M. J., Heyland A., Hodin J., Brandhorst B. P., 2006 Interspecific variation in metamorphic competence in marine invertebrates: the significance for comparative investigations into the timing of metamorphosis. Integrative and Comparative Biology 46: 662–682.

Bonner J. T., 2003 Evolution of development in the cellular slime molds. Evolution & development 5: 305–313.

Breakspear A., Momany M., 2007 Aspergillus nidulans conidiation genes dewA, fluG, and stuA are differentially regulated in early vegetative growth. Eukaryotic cell 6: 1697–1700.

Browse J., 2009 Jasmonate Passes Muster: A Receptor and Targets for the Defense Hormone. Annual Review of Plant Biology 60: 183–205.

Butnick N. Z., Yager L. N., Kurtz M. B., Champe S. P., 1984 Genetic analysis of mutants of Aspergillus nidulans blocked at an early stage of sporulation. Journal of Bacteriology 160: 541–545.

Calcagno-Pizarelli A. M., Negrete-Urtasun S., Denison S. H., Rudnicka J. D., Bussink H. J., Munera-Huertas T., Stanton L., Hervas-Aguilar A., Espeso E. A., Tilburn J., Arst H. N., Peñalva M. A., 2007 Establishment of the Ambient pH Signaling Complex in Aspergillus nidulans: PalI Assists Plasma Membrane Localization of PalH. Eukaryotic cell 6: 2365–2375.

Calo S., Shertz-Wall C., Lee S. C., Bastidas R. J., Nicolás F. E., Granek J. A., Mieczkowski P., Torres-Martínez S., Ruiz-Vázquez R. M., Cardenas M. E., Heitman J., 2014 Antifungal drug resistance evoked via RNAi-dependent epimutations. Nature 513: 555–558.

Casselton L., Zolan M., 2002 The art and design of genetic screens: filamentous fungi. Nature Reviews Genetics 3: 683–697.

Champe S. P., Kurtz M. B., Yager L. N., Butnick N., Axelrod D. E., 1981 Spore formation in Aspergillus nidulans: competence and other developmental processes.

Champe S., Nagle D., Yager L. N., 1994 Sexual sporulation. Progress in industrial Microbiology 29: 429–454.

Chen K., Wallis J. W., McLellan M. D., Larson D. E., Kalicki J. M., Pohl C. S., McGrath S. D., Wendl M. C., Zhang Q., Locke D. P., Shi X., Fulton R. S., Ley T. J., Wilson R. K., Ding L., Mardis E. R., 2009 BreakDancer: an algorithm for high-resolution mapping of genomic structural variation. Nature methods 6: 677–681.

Cingolani P., Platts A., Wang L. L., Coon M., Nguyen T., Wang L., Land S. J., Lu X., Ruden D. M., 2014 A program for annotating and predicting the effects of single nucleotide polymorphisms, SnpEff. Fly 6: 80–92.

Clutterbuck A. J., 1969 A mutational analysis of conidial development in Aspergillus nidulans. Genetics 63: 317–327.

Dynesen J., Nielsen J., 2003 Branching is coordinated with mitosis in growing hyphae of Aspergillus nidulans. Fungal Genetics and Biology 40: 15–24.

Etxebeste O., Ni M., Garzia A., Kwon N.-J., Fischer R., Yu J. H., Espeso E. A., Ugalde U., 2008 Basic-Zipper-Type Transcription Factor FlbB Controls Asexual Development in Aspergillus nidulans. Eukaryotic cell 7: 38–48.

Etxebeste O., Herrero-García E., Araújo-Bazan L., Rodríguez-Urra A. B., Garzia A., Ugalde U., Espeso E. A., 2009 The bZIP-type transcription factor FlbB regulates distinct morphogenetic stages of colony formation in Aspergillus nidulans. Molecular Microbiology 73: 775–789.

Etxebeste O., Garzia A., Espeso E. A., Ugalde U., 2010 Aspergillus nidulans asexual development: making the most of cellular modules. Trends in Microbiology 18: 569–576.

Evans J. L., Gealt M. A., 1985 The sterols of growth and stationary phases of Aspergillus nidulans cultures. Journal of general microbiology 131: 279–284.

Fiddy C., Fiddy C., Trinci A. P., 1976 Mitosis, septation, branching and the duplication cycle in Aspergillus nidulans. Journal of general microbiology 97: 169–184.

Fox E. M., Howlett B. J., 2008 Secondary metabolism: regulation and role in fungal biology. Current opinion in microbiology 11: 481–487.

Garzia A., Etxebeste O., Herrero-García E., Fischer R., Espeso E. A., Ugalde U., 2009 Aspergillus nidulansFlbE is an upstream developmental activator of conidiation functionally associated with the putative transcription factor FlbB. Molecular Microbiology 71: 172–184.

Gealt M. A., Axelrod D. E., 1974 Coordinate regulation of enzyme inducibility and developmental competence in Aspergillus nidulans. Developmental Biology 41: 224–232.

Gilchrist M. A., Sulsky D. L., Pringle A., 2006 Identifying fitness and optimal life-history strategies for an asexual filamentous fungus. Evolution 60: 970–979.

Gluckman P. D., Hanson M. A., 2006 Evolution, development and timing of puberty. Trends in endocrinology and metabolism: TEM 17: 7–12.

Goldman G. H., Osmani S. A. (Eds.), 2007 The Aspergilli.

Gomori G., 1946 Buffers in the range of pH 6.5 to 9.6. Proceedings of the Society for Experimental Biology and Medicine. Society for Experimental Biology and Medicine (New York, N.Y.) 62: 33.

Gravelat F. N., Doedt T., Chiang L. Y., Liu H., Filler S. G., Patterson T. F., Sheppard D. C., 2008 In Vivo Analysis of Aspergillus fumigatus Developmental Gene Expression Determined by Real-Time Reverse Transcription-PCR. Infection and immunity 76: 3632–3639.

Gressel J., Gressel J., Galun E., Galun E., 1967 Morphogenesis in Trichoderma: photoinduction and RNA. Developmental Biology 15: 575–598.

Hadley G., Harrold C. E., 1958 The sporulation of Penicillium notatum Westling in submerged liquid culture. II. The initial sporulation phase. J. Exp. Bot 9: 418–425.

Hall N. E., Axelrod D. E., 1977 Interference of cellular ferric ions with DNA extraction and the application to methods of DNA determination. Analytical biochemistry 79: 425–430.

Hall N. E., Axelrod D. E., 1978 Sporulation competence in Aspergillus nidulans: a role for iron in development. Cell differentiation 7: 73–82.

Han K. H., Han K. Y., Yu J. H., Chae K. S., Jahng K. Y., Jahng K. Y., Han D. M., 2001 The nsdD gene encodes a putative GATA-type transcription factor necessary for sexual development of Aspergillus nidulans. Molecular Microbiology 41: 299–309.

Han K.-H., 2009 Molecular Genetics of Emericella nidulans Sexual Development. Mycobiology 37: 171–182.

Harispe L., Portela C., Scazzocchio C., Peñalva M. A., Gorfinkiel L., 2008 Ras GTPase-Activating Protein Regulation of Actin Cytoskeleton and Hyphal Polarity in Aspergillus nidulans. Eukaryotic cell 7: 141–153.

Heaton L. L. M., Jones N. S., Fricker M. D., 2016 Energetic Constraints on Fungal Growth. The American Naturalist 187: E27–E40.

Hind S. R., Pulliam S. E., Veronese P., Shantharaj D., Nazir A., Jacobs N. S., Stratmann J. W., 2011 The COP9 signalosome controls jasmonic acid synthesis and plant responses to herbivory and pathogens. The Plant Journal 65: 480–491.

Hynes M. J., Kelly J. M., 1977 Pleiotropic mutants of Aspergillus nidulans altered in carbon metabolism. Molecular & general genetics: MGG 150: 193–204.

Istock C. A., Istock C. A., 1967 Evolution of Complex Life Cycle Phenomena - an Ecological Perspective. Evolution; international journal of organic evolution 21: 592–605.

Johnston G. C., Pringle J. R., Hartwell L. H., 1977 Coordination of growth with cell division in the yeast Saccharomyces cerevisiae. Experimental Cell Research 105: 79–98.

Kawasaki L., Aguirre J., 2001 Multiple catalase genes are differentially regulated in Aspergillus nidulans. Journal of Bacteriology 183: 1434–1440.

Keller N. P., Turner G., Bennett J. W., 2005 Fungal secondary metabolism: from biochemistry to genomics. Nature Reviews Microbiology 3: 937–947.

Kelley D. R., Schatz M. C., Salzberg S. L., 2010 Quake: quality-aware detection and correction of sequencing errors. Genome Biology 11: R116.

Kennedy P. A., 1999 The velvet gene encodes a novel signal integrator that controls the initiation of conidiation in Aspergillus nidulans.

Kent W. J., Baertsch R., Hinrichs A., Miller W., Haussler D., 2011 Evolution’s cauldron: Duplication, deletion, and rearrangement in the mouse and human genomes. Proceedings of the National Academy of Sciences 100: 11484–11489.

Kurtz M. B., Champe S. P., 1979 Genetic control of transport loss during development of Aspergillus nidulans. Developmental Biology.

Kurtz M. B., 1980 Regulation of Fructose Transport During Growth of Aspergillus nidulans. Microbiology 118: 389–396.

Kurtz S., Phillippy A., Delcher A. L., Smoot M., Shumway M., Antonescu C., Salzberg S. L., 2004 Versatile and open software for comparing large genomes. Genome Biology 5: R12.

Kwon N.-J., Garzia A., Espeso E. A., Ugalde U., Yu J.-H., 2010 FlbC is a putative nuclear C2H2 transcription factor regulating development in Aspergillus nidulans. Molecular Microbiology 77: 1203–1219.

Lee M. V., Topper S. E., Hubler S. L., Hose J., Wenger C. D., Coon J. J., Gasch A. P., 2011 A dynamic model of proteome changes reveals new roles for transcript alteration in yeast. Molecular Systems Biology 7: 514–514.

Leeuwen M. R. van, Krijgsheld P., Bleichrodt R., Menke H., Stam H., Stark J., Wosten H. A. B., Dijksterhuis J., 2013 Germination of conidia of Aspergillus niger is accompanied by major changes in RNA profiles. Studies in mycology 74: 59–70.

Li H., Handsaker B., Wysoker A., Fennell T., Ruan J., Homer N., Marth G., Abecasis G., Durbin R., 1000 Genome Project Data Processing Subgroup, 2009 The Sequence Alignment/Map format and SAMtools. Bioinformatics 25: 2078–2079.

Li H., Durbin R., 2010 Fast and accurate long-read alignment with Burrows-Wheeler transform. Bioinformatics 26: 589–595.

Li H., 2011 A statistical framework for SNP calling, mutation discovery, association mapping and population genetical parameter estimation from sequencing data. Bioinformatics 27: 2987–2993.

Maere S., Heymans K., Kuiper M., 2005 BiNGO: a Cytoscape plugin to assess overrepresentation of Gene Ontology categories in Biological Networks. Bioinformatics 21: 3448–3449.

Marin F. T., 1976 Regulation of development in Dictyostelium discoideum: I. Initiation of the growth to development transition by amino acid starvation. Developmental Biology 48: 110–117.

McKenna A., Hanna M., Banks E., Sivachenko A., Cibulskis K., Kernytsky A., Garimella K., Altshuler D., Gabriel S., Daly M., DePristo M. A., 2010 The Genome Analysis Toolkit: A MapReduce framework for analyzing next-generation DNA sequencing data. Genome Research 20: 1297–1303.

Miller K. Y., Toennis T. M., Toennis T. M., Adams T. H., Miller B. L., 1991 Isolation and transcriptional characterization of a morphological modifier: the Aspergillus nidulans stunted (stuA) gene. Molecular & general genetics: MGG 227: 285–292.

Mooney J. L., Yager L. N., 1990 Light is required for conidiation in Aspergillus nidulans. Genes & Development 4: 1473–1482.

Moran N. A., 1994 Adaptation and Constraint in the Complex Life-Cycles of Animals. Annual Review of Ecology and Systematics 25: 573–600.

Morton A. G., England D. J. F., Towler D. A., 1958 The physiology of sporulation in Penicillium griseofulvum Dierckx. Transactions of the British Mycological Society 41: 39–51.

Nahlik K., Dumkow M., Bayram O., Helmstaedt K., Busch S., Valerius O., Gerke J., Hoppert M., Schwier E., Opitz L., Westermann M., Grond S., Feussner K., Goebel C., Kaever A., Meinicke P., Feussner I., Braus G. H., 2010 The COP9 signalosome mediates transcriptional and metabolic response to hormones, oxidative stress protection and cell wall rearrangement during fungal development. Molecular Microbiology 78: 964–979.

Nayak T., 2005 A Versatile and Efficient Gene-Targeting System for Aspergillus nidulans. Genetics 172: 1557–1566.

Negrete-Urtasun S., Negrete-Urtasun S., Reiter W., Reiter W., Diez E., Diez E., Denison S. H., Denison S. H., Tilburn J., Tilburn J., Espeso E. A., Peñalva M. A., Peñalva M. A., Arst H. N., 1999 Ambient pH signal transduction in Aspergillus: completion of gene characterization. Molecular Microbiology 33: 994–1003.

Noble L. M., Andrianopoulos A., 2013 Reproductive competence: a recurrent logic module in eukaryotic development. Proceedings of the Royal Society B: Biological Sciences 280: 20130819–20130819.

Oshlack A., Emslie D., Corcoran L., Smyth G. K., 2007 Normalization of boutique two-color microarrays with a high proportion of differentially expressed probes. Genome Biology 8: R2.

Pastushok M., Pastushok M., Axelrod D. E., 1976 Effect of glucose, ammonium and media maintenance on the time of conidiophore initiation by surface colonies of Aspergillus nidulans. Journal of general microbiology 94: 221–224.

Pearse A. G. E., 1991 Histochemistry, Theoretical and Applied: Enzyme histochemistry.

Poethig R. S., 2003 Phase change and the regulation of developmental timing in plants. Science 301: 334–336.

Poethig R. S., 2010 The past, present, and future of vegetative phase change. Plant Physiology 154: 541–544.

Pontecorvo G., 1952 Genetic analysis without sexual reproduction by means of polyploidy in Aspergillus niger. The Journal of General Microbiology.

Prade R. A., Timberlake W. E., 1993 The Aspergillus nidulans brlA regulatory locus consists of overlapping transcription units that are individually required for conidiophore development. The EMBO Journal 12: 2439–2447.

Ritchie M. E., Silver J., Oshlack A., Holmes M., Diyagama D., Holloway A., Smyth G. K., 2007 A comparison of background correction methods for two-colour microarrays. Bioinformatics 23: 2700–2707.

Rittenour W. R., Si H., Harris S. D., 2009 Hyphal morphogenesis in Aspergillus nidulans. Fungal Biology Reviews 23: 20–29.

Robson G., Robson G., Prebble E., Prebble E., Rickers A., Rickers A., Hosking S., Hosking S., Denning D., Denning D., Trinci A., Robertson W., Robertson W., 1996 Polarized Growth of Fungal Hyphae Is Defined by an Alkaline pH Gradient. Fungal genetics and biology: FG & B 20: 289–298.

Rodriguez-Romero J., Hedtke M., Kastner C., Müller S., Fischer R., 2010 Fungi, Hidden in Soil or Up in the Air: Light Makes a Difference. Annual Review of Microbiology 64: 585–610.

Rodriguez-Urra A. B., Jimenez C., Nieto M. I., Rodriguez J., Hayashi H., Ugalde U., 2012 Signaling the Induction of Sporulation Involves the Interaction of Two Secondary Metabolites in Aspergillus nidulans. ACS chemical biology 7: 599–606.

Roncal T., Roncal T., Ugalde U. O., Irastorza A., Irastorza A., 1993 Calcium-induced conidiation in Penicillium cyclopium: calcium triggers cytosolic alkalinization at the hyphal tip. Journal of Bacteriology 175: 879–886.

Roncal T., Cordobes S., Sterner O., Ugalde U., 2002 Conidiation in Penicillium cyclopium Is Induced by Conidiogenone, an Endogenous Diterpene. Eukaryotic cell 1: 823–829.

Roncal T., Ugalde U., 2003 Conidiation induction in Penicillium. Research in Microbiology 154: 539–546.

Roper M., Ellison C., Taylor J. W., Glass N. L., 2011 Nuclear and Genome Dynamics in Multinucleate Ascomycete Fungi. Current Biology 21: R786–R793.

Ruger-Herreros C., Rodriguez-Romero J., Fernandez-Barranco R., Olmedo M., Fischer R., Corrochano L. M., Canovas D., 2011 Regulation of Conidiation by Light in Aspergillus nidulans. Genetics 188: 809–822.

Semighini C. P., Harris S. D., 2008 Regulation of Apical Dominance in Aspergillus nidulans Hyphae by Reactive Oxygen Species. Genetics 179: 1919–1932.

Sewall T. C., Mims C. W., Timberlake W. E., 1990 abaA controls phialide differentiation in Aspergillus nidulans. The Plant cell 2: 731–739.

Sheppard D. C., 2005 The Aspergillus fumigatus StuA Protein Governs the Up-Regulation of a Discrete Transcriptional Program during the Acquisition of Developmental Competence. Molecular biology of the cell 16: 5866–5879.

Simpson J. T., Wong K., Jackman S. D., Schein J. E., Jones S. J. M., Birol I., 2009 ABySS: a parallel assembler for short read sequence data. Genome Research 19: 1117–1123.

Smyth G. K., Speed T., 2003 Normalization of cDNA microarray data. Methods (San Diego, Calif) 31: 265–273.

Smyth G. K., 2004 Linear Models and Empirical Bayes Methods for Assessing Differential Expression in Microarray Experiments. Statistical applications in genetics and molecular biology 3: 1–25.

Smyth G. K., Michaud J., Scott H. S., 2005 Use of within-array replicate spots for assessing differential expression in microarray experiments. Bioinformatics 21: 2067–2075.

Soid-Raggi G., Sánchez O., Aguirre J., 2006 TmpA, a member of a novel family of putative membrane flavoproteins, regulates asexual development in Aspergillus nidulans. Molecular Microbiology 59: 854–869.

Som T., Kolaparthi V. S., 1994 Developmental decisions in Aspergillus nidulans are modulated by Ras activity. Molecular and Cellular Biology 14: 5333–5348.

Stoodley P., Sauer K., Davies D. G., Costerton J. W., 2002 Biofilms as Complex Differentiated Communities. Annual Review of Microbiology 56: 187–209.

Sutton K. W., 1999 Analysis of acoC: a gene involved in developmental commitment in Aspergillus nidulans.

Todd R. B., Davis M. A., Hynes M. J., 2007a Genetic manipulation of Aspergillus nidulans: meiotic progeny for genetic analysis and strain construction. Nature Protocols 2: 811–821.

Todd R. B., Davis M. A., Hynes M. J., 2007b Genetic manipulation of Aspergillus nidulans: heterokaryons and diploids for dominance, complementation and haploidization analyses. Nature Protocols 2: 822–830.

Trinci A. P., 1974 A study of the kinetics of hyphal extension and branch initiation of fungal mycelia. Journal of general microbiology 81: 225–236.

Truman J. W., Riddiford L. M., 1999 The origins of insect metamorphosis. Nature 401: 447–452.

Tsitsigiannis D. I., 2004 The Lipid Body Protein, PpoA, Coordinates Sexual and Asexual Sporulation in Aspergillus nidulans. Journal of Biological Chemistry 279: 11344–11353.

Turner J. J., Ewald J. C., Skotheim J. M., 2012 Cell size control in yeast. Current biology: CB 22: R350–9.

Vanetten J. L., Gottlieb D., 1965 Biochemical changes during the growth of fungi. II. Ergosterol and fatty acids in Penicillium atrovenetum. Journal of Bacteriology 89: 409–414.

Vining L., 1990 Functions of secondary metabolites. Annual Reviews in Microbiology.

Vogel C., Silva G. M., Marcotte E. M., 2011 Protein Expression Regulation under Oxidative Stress. Molecular & Cellular Proteomics 10: M111.009217–M111.009217.

Vogel C., Marcotte E. M., 2012 Insights into the regulation of protein abundance from proteomic and transcriptomic analyses. Nature Reviews Genetics 13: 227–232.

Wiegmann B. M., Trautwein M. D., Kim J.-W., Cassel B. K., Bertone M. A., Winterton S. L., Yeates D. K., 2009 Single-copy nuclear genes resolve the phylogeny of the holometabolous insects. BMC biology 7: 34–640.

Wilbur H. M., 1980 Complex Life-Cycles. Annual Review of Ecology and Systematics 11: 67–93.

Wong K. H., Hynes M. J., Davis M. A., 2008 Recent Advances in Nitrogen Regulation: a Comparison between Saccharomyces cerevisiae and Filamentous Fungi. Eukaryotic cell 7: 917–925.

Yager L., Kurtz M., 1982 Temperature-shift analysis of conidial development in Aspergillus nidulans. Developmental Biology: 1–12.

Ye K., Schulz M. H., Long Q., Apweiler R., Ning Z., 2009 Pindel: a pattern growth approach to detect break points of large deletions and medium sized insertions from paired-end short reads. Bioinformatics 25: 2865–2871.

Young R., 1976 Fat, energy and mammalian survival. American Zoologist.

Yu J.-H., Keller N., 2005 Regulation of secondary metabolism in filamentous fungi. Annual review of phytopathology 43: 437–458.

Zacharias L., Zacharias L., 1969 Age at menarche. New England Journal of Medicine.

Zemla A., Thomas Y., Kedziora S., Knebel A., Wood N. T., Rabut G., Kurz T., 2013 CSN-and CAND1-dependent remodelling of the budding yeast SCF complex. Nature communications 4: 1641.

